# Encoding of predictive associations in human prefrontal and medial temporal neurons during Pavlovian conditioning

**DOI:** 10.1101/2023.02.10.528055

**Authors:** Tomas G. Aquino, Hristos Courellis, Adam N. Mamelak, Ueli Rutishauser, John P. O’Doherty

## Abstract

Pavlovian conditioning is thought to involve the formation of learned associations between stimuli and values, and between stimuli and specific features of outcomes. Here we leveraged human single neuron recordings in ventromedial prefrontal, dorsomedial frontal, hippocampus and amygdala neurons while patients performed a sequential Pavlovian conditioning task containing both stimulus-value and stimulus-stimulus associations. Neurons in the ventromedial prefrontal cortex encoded predictive value along with the amygdala, but also encoded predictions about the identity of stimuli that would subsequently be presented, suggesting a role for neurons in this region in encoding predictive information beyond value. Unsigned error signals were found in dorsomedial prefrontal areas and hippocampus, potentially supporting learning of non-value related outcome features. Our findings implicate distinct human prefrontal and medial temporal neuronal populations in mediating predictive associations which could partially support model-based mechanisms during Pavlovian conditioning.

**Significance statement:** Pavlovian conditioning is a fundamental form of learning, allowing organisms to associate stimuli and outcomes. Recent Pavlovian work suggests that phenomena such as devaluation sensitivity and sensory preconditioning can be explained by a model-based learning framework. How human neurons perform model-based learning during Pavlovian conditioning is still an open question. We recorded single neurons from epilepsy patients during a two-step Pavlovian conditioning task and found that ventromedial prefrontal neurons encoded expected rewards along with amygdala neurons, but also predicted the identity of upcoming stimuli as required for model-based cognition. Additionally, medial frontal neurons were found to encode error signals that could be used for stimulus-outcome learning. This is the first study mapping model-based computations during Pavlovian conditioning in human neurons.

## 1 Introduction

In Pavlovian conditioning, an animal learns to make predictions about future affectively significant events such as an appetitive outcome so as to facilitate prospective behavioral adaptations in the form of conditioned responses.^1, 2^ Elucidating the nature of the associations that underpin the acquisition of Pavlovian conditioned responses, and of the computational mechanism by which such associations are learned, has long been an active area of research at both behavioral and neural levels.^2–6^

Perhaps the most influential theory of Pavlovian conditioning is the Rescorla-Wagner (RW) rule and variants thereof. This class of model assumes that conditioning occurs via a prediction error (PE) that reports on discrepancies between expectations associated with the conditioned stimulus (CS), and the unconditioned stimulus (US) or experienced outcome.^7^ This PE is used to update associative weights, driving the degree of elicitation of a conditioned response (CR) by the CS. Such a model can account for a wide variety of conditioning phenomena observed in behavioral experiments such as blocking, over-expectation and conditioned inhibition.^8–10^ Compelling empirical support for the role of RW-type PEs in appetitive Pavlovian learning has emerged from the finding that the phasic activity of dopamine neurons in the midbrain resembles a reward-specific prediction error, manifesting many of the specific predictions of the RW model,^11–13^ while also being causally relevant for driving learning of appetitive Pavlovian conditioned responses.^14–16^

However, the RW model has long been known to fail to account for numerous phenomena observed in Pavlovian conditioning, such as sensory preconditioning^17, 18^ and resistance of inhibitory stimuli to extinction.^19^ Moreover, many Pavlovian conditioned responses are known to be devaluation or revaluation sensitive, in which changes in the value of an unconditioned stimulus can produce immediate changes in the conditioned responses elicited to a corresponding CS,^20^ inconsistent with the trial and error updating of conditioned responses learned via reward PEs. To underpin such behavioral phenomena, other forms of associations have been suggested to be formed in the brain, including stimulus-stimulus associations.^1, 21^ These types of stimulus-stimulus associations can be viewed as one of the constituent parts of “model-based” Pavlovian learning, in which features about an outcome such as predictions about its identity are used to drive on-line updating about value analogous to the model-based algorithms thought to be important in instrumental conditioning.^22–25^

Consistent with this richer associative framework, accumulating evidence suggests that the brain encodes associative predictions about stimuli beyond just value. In particular, activity in orbitofrontal cortex neurons in rodents has been found to signal changes in stimulus features such as stimulus identity.^26^ fMRI studies in humans also found evidence for encoding of predictive outcome identity representations during reward learning in OFC and anterior dorsal striatum.^24, 25, 27, 28^

Learning about stimulus features other than value is theoretically proposed to be mediated by a kind of prediction error signal known as the state prediction error or identity prediction error.^23, 24, 28–31^ This type of error signal is distinct from reward prediction error, and encodes discrepancies between an agent’s expectation about which stimulus will occur and which stimulus actually occurs, irrespective of the amount of reward or value involved. Such error signals have been found to be present in many brain areas in human fMRI studies, including a frontoparietal network encompassing dorsomedial prefrontal cortex, as well as in dopaminergic regions of the midbrain, the OFC and striatum.^23, 30, 32–35^ In rodents, dopamine neurons themselves have been suggested to be sensitive to violations in predicted identity beyond value, and are even found to be necessary for the acquisition of learned predictions about stimulus identity.^29, 36^

The goal of the present study was to investigate neural representations of multiple forms of Pavlovian associative learning at the cellular level in humans. A key limitation of previous studies on the neural mechanisms of Pavlovian conditioning in humans to date has been that these studies have relied on BOLD fMRI. While this technique can provide a bird’s eye view on the neural representations present in different brain regions, its lack of spatial and temporal resolution precludes insight into the underlying neuronal representations and computations taking place during Pavlovian learning. Probing activity at the level of single neurons is of fundamental importance both for understanding how Pavlovian conditioning is implemented in the human brain, and for gaining insights at the cellular level so that they can be compared to the large body of single-neuron work done in animal model systems.

To accomplish this goal, we conducted single neuron recordings in patients undergoing intracranial monitoring as part of the neurosurgical treatment for refractory epilepsy. Patients performed a Pavlovian learning task in which pairs of visual stimuli (fractals) were presented in a sequence that was associated with either the subsequent delivery of a reward outcome (a picture of a food item that could subsequently be consumed) or a non-reward (no outcome). Across blocks, the particular stimuli associated with each outcome were changed to dissociate predictive coding related to the identity of a stimulus from coding of the predicted reward-value of the outcome associated with that stimulus. Pairs of stimuli were presented in a sequence to test whether neurons encoded predictions about the identity of the stimulus expected to occur next in that sequence. This design allowed for differentiating between neurons encoding stimulus-stimulus relationships and stimulus-reward associations. In addition, this paradigm afforded the opportunity to assess whether there exist neurons that encode prediction error signals beyond the canonical reward prediction error, which could potentially play a role in model-based learning.

We obtained simultaneous recordings from neurons in the ventromedial prefrontal cortex (including neurons on the medial orbital surface) as well as the amygdala, two brain areas that have long been suggested to play a role in value-based learning and Pavlovian learning in particular.^25, 37–46^ We also obtained recordings from the hippocampus, a brain region that has been previously implicated in model-based inference,^28, 47, 48^ as well as from the anterior cingulate cortex and pre-supplementary motor cortex (preSMA) in dorsomedial frontal cortex, a region that has been previously found to prominently encode error signals.^49–53^

We hypothesized that neurons in ventromedial prefrontal cortex and amygdala might play a role in encoding predictive information about stimulus identity, consistent with a role for these structures in stimulus-stimulus learning, and in model-based inference more generally.^18, 24, 25, 28^ In addition to testing for neurons correlating with stimulus identity we also tested for neurons encoding stimulus-value associations. Furthermore, we tested whether neural correlates of state prediction errors that could underpin learning of stimulus-stimulus associations in a model-based learning mechanism for Pavlovian conditioning could be found at the single neuron level in any of the recorded brain structures. We then sought to determine whether such neuronal signals could be clearly dissociated from reward prediction errors or outcome signals. Finally, the fact that we could record from multiple structures simultaneously provided us with the opportunity to gain insight into the network-level interactions between neurons in these structures as a function of learning.

## 2 Results

### 2.1 Behavioral evidence of Pavlovian conditioning

We recorded 165 AMY, 119 HIP, 86 vmPFC, 137 preSMA, and 103 dACC single neurons (610 total) in 13 sessions from 12 patients implanted with hybrid macro/micro electrodes for epilepsy monitoring (Fig. 1C). Patients performed a sequential Pavlovian conditioning task (Fig. 1A) with two conditioned stimuli in the form of fractal images: distal (CSd), followed by proximal (CSp). Conditioned stimuli were then followed by an outcome, which could be rewarding or neutral.^25, 54^ Outcomes were delivered in the form of videos, either of a hand depositing a piece of candy in a bag, or of an empty hand approaching a bag. Patients were told that every display of the rewarding video contributed partially to the amount of real candy they would be given after the end of the session. Patients were asked to pay attention to CS identities as they would be predictive of rewards. Subjects were asked to indicate by button which side of the screen the CSp stimulus was shown to verify attention, but were informed that their accuracy or speed of doing so did not influence the outcome of the trial in any way. In each of the four blocks, a CSd/CSp pair would be more likely associated with the reward, and were therefore defined as CSd+/CSp+ (see Materials and Methods for task details), according to a common/rare transition structure (Fig. 1D). Conversely, the other CSd/CSp pair was more likely to precede the neutral outcome, and are referred to as CSdn/CSpn.

**Figure 1:**
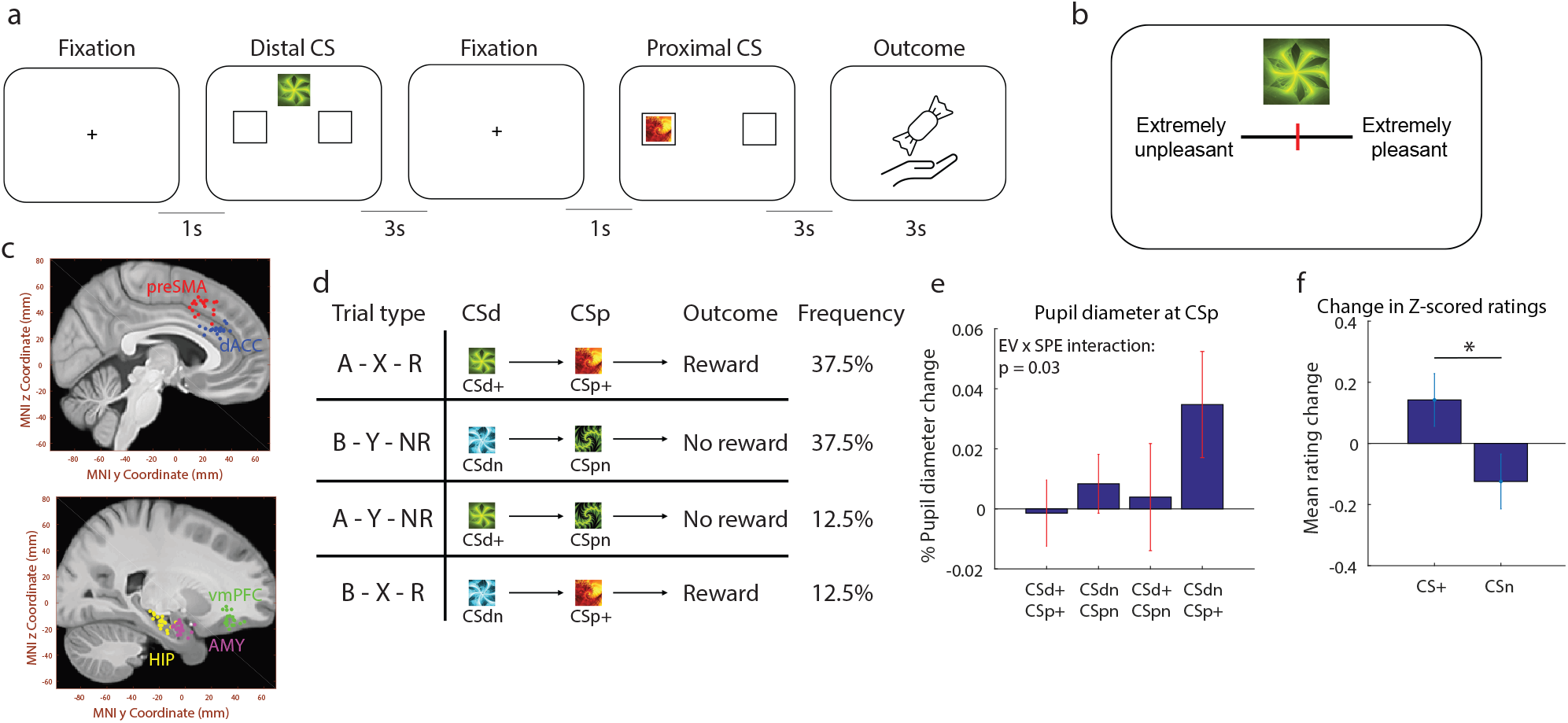
Pavlovian conditioning task and behavior. (a) Trial structure. After a fixation period, patients saw a sequence of two conditioned stimuli, distal and proximal, with a 1s fixation period in between them. Then, outcomes were presented: for positive outcomes, a video of a hand depositing a piece of candy in a bag; for neutral outcomes, an empty hand approaching a bag. (b) Rating screen. Before the task and between blocks, patients used a sliding bar to rate fractal stimuli according to their subjective preference, from extremely unpleasant to extremely pleasant. (c) Electrode positioning. Each dot indicates the location of a microwire bundle in amygdala (magenta), hippocampus (yellow), preSMA (red), dACC (blue), or vmPFC (green). (d) Trial types. Stimuli transitioned from distal to proximal according to a common/rare probabilistic structure. The same 2 fractals were used as CSp throughout the entire task, while new fractals were picked as CSd in every block. (e) Pupil diameter change at CSp. We compared pupil diameter during CSp presentation with a baseline period, for each trial type. Error bars represent SEM. (f) Changes in stimulus ratings. After each block, patients rated stimuli for their subjective preference. We compared how ratings changed for each fractal compared to its previous value, depending on whether they were a positive or neutral CS in that block.

To characterize value learning and stimulus predictions in this task, we fit a normative model-based transition matrix model (see Materials and Methods for model details) to infer expected values (EV), state prediction errors (SPE) and transition probabilities on a trial by trial basis, for each session (Fig. 1E). With these transition probabilities, we could also estimate which CSp was most likely to follow a CSd in each trial, which we refer to as *CSp presumed identity*. To infer whether Pavlovian conditioning occurred across patients, we correlated the obtained model covariates with behavioral metrics indicative of conditioning: stimulus ratings and pupil dilation.

Subjective preference ratings for all fractal images were obtained in the beginning of the task. Additionally, between blocks, we asked patients to re-rate the fractals that were included in the previous block to obtain measures of changes in subjective preference as a function of patients’ experience in the task (Fig. 1B). When grouping CSd and CSp together, we observed that stimuli used as CS+ were rated significantly higher than stimuli used as CSn (p = 0.02, one sided t-test, Fig. 1F). We also tested whether distal and proximal stimuli had their ratings change by a different amount by contrasting absolute rating changes for (CSd+,CSdn) versus (CSp+,CSpn), and found no significant difference for distal vs. proximal stimuli (p=0.16, two-tailed t-test). Finally, we found that initial ratings for stimuli which would be used as CSd+ and CSdn did not significantly differ (*p* = 0.91, t-test), further indicating stimulus rating changes were a consequence of experiencing the task.

We performed eye tracking simultaneously with single-neuron recordings to further assess the success of conditioning (see methods). Pupil diameter was analyzed in two distinct time windows: during CSd presentation and CSp presentation (see Materials and Methods for details on pupil analysis). We obtained the average pupil diameter change within these periods, relative to a baseline, and tested whether they correlated with model covariates with a linear mixed effects model, with session number as a random effect. Specifically, the model for pupil diameter during CSd presentation included the EV of the CSd, while the model for pupil diameter during CSp presentation included CSp EV, SPE, and an interaction term between CSp EV and SPE. We found no effect for CSd EV in the first model (p = 0.30). In the second model (Fig. 1E), there was no significant effect of CSp EV (p = 0.07) nor SPE (p = 0.06), but there was a significant interaction between CSp EV and SPE (p = 0.02). This result indicates that pupil diameter correlated with a combination of computational factors inferred from the model-based framework. A similar interaction was previously observed in a Pavlovian conditioning paradigm performed in a neurotypical population.^54^

Overall, the aggregate behavioral evidence from changes in subjective stimulus ratings and pupil sensitivity to an interaction of EV and SPE indicates collective evidence of Pavlovian conditioning across our subject sample.

### 2.2 Predictive identity coding in vmPFC

We next investigated whether firing rates in individual neurons correlated with task variables and the estimated computational model-derived covariates using a Poisson GLM analysis (see Materials and Methods for details). We obtained spike counts for each neuron in the time windows that were relevant for each regressed variable (e.g. counting spikes during outcome presentation for regressing outcomes). After obtaining the number of neurons significant for a given criterion, we tested whether the number of significant neurons in each brain region was more than expected by chance with a binomial test (Bonferroni corrected for the number of tested brain areas).

We first regressed the CSp presumed identity (i.e. the most likely proximal stimulus identity) with the firing rate of neurons at the time of CSd presentation (Fig. 2A) to test whether the most likely identity of the next presented stimulus was already encoded by neurons at distal time. 11.5% of vmPFC neurons encoded CSp presumed identity (*p* < 0.05, binomial test), indicating that vmPFC neurons represent predicted stimulus identity in a stimulus-stimulus association context. On the other hand, we found that no brain areas significantly encoded the actual identity of the proximal stimulus at the time of proximal stimulus presentation (*p* > 0.05 for all regions, binomial test).

**Figure 2:**
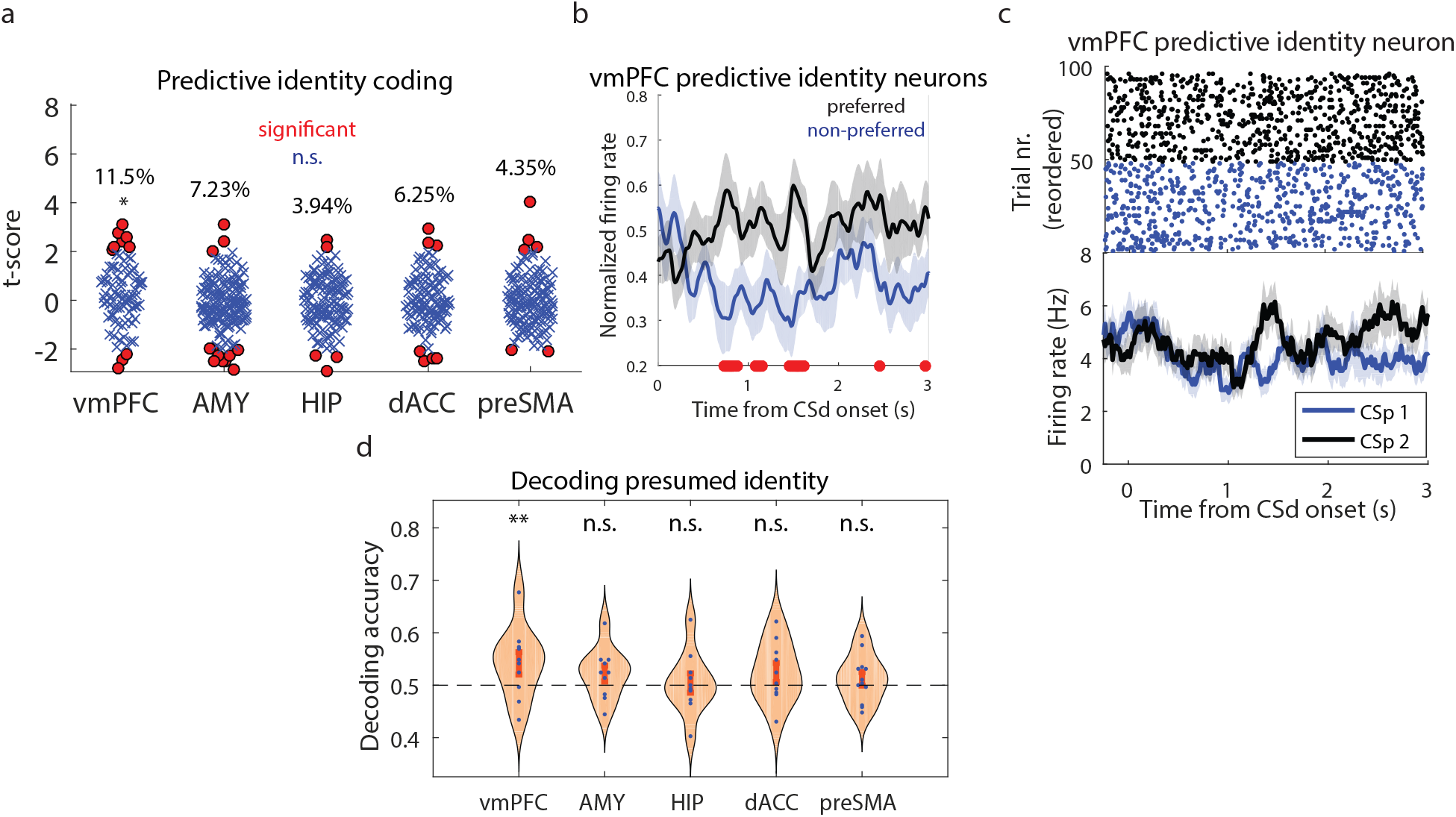
Predictive identity coding in vmPFC. (a) T-scores for every neuron in each brain area, for a GLM predicting spike counts during CSd presentation with the presumed identity of the CSp as a regressor. Red dots indicate significant neurons and stars indicate significance across the entire region, corrected across areas. (b) Time course of normalized firing rates in vmPFC neurons which coded stimulus identities predictively. Trials are separated between preferred (black) versus non-preferred (blue) stimulus identities, and red dots indicate time points with a significant difference between trajectories. (c) vmPFC neuron whose activity during distal presentation correlates with the presumed identity of the proximal stimulus (blue: first CSp; black: second CSp). Top: raster plot; Bottom: PSTH. (d) Decoding accuracy for presumed proximal identity during distal presentation. Each dot indicates accuracy in one session, stars indicate significance across sessions with a bootstrapped null distribution, corrected across areas. Bars and dashed lines indicate standard error and chance level, respectively.

For vmPFC neurons whose response signaled predictive identity coding, we summarized their normalized firing rates over time, separated by trials containing their preferred versus non-preferred identity (Fig. 2B). For each time point, we tested whether preferred versus non-preferred firing rates were different, and found that they first differed 0.73s after CSd onset (*p* < 0.05, t-test, uncorrected). An example neuron performing this type of encoding is shown in Fig. 2C. We also regressed the EV of the presumed CSp at distal time and the actual CSp identity at proximal time with firing rate but did not find a significant neuron count in any region (*p* > 0.05 for all regions, binomial test).

We next turned to population level analysis to investigate if the joint activity patterns from all neurons recorded simultaneously in a given brain area were predictive of the variables of interest. All population level decoding analysis throughout this paper is performed session-by-session, only utilizing neurons that were recorded simultaneously. We performed cross-validated population decoding analysis with a linear SVM, obtaining significance levels with a bootstrapped null distribution (see Materials and Methods for details). We found that CSp presumed identity could be significantly decoded at distal time in vmPFC (*p* < 0.01, permutation test), in consonance with the single neuron selection result discussed above (Fig. 2D). These results further establish vmPFC neural activity as a substrate for predictive coding during learning of stimulus-stimulus associations in Pavlovian conditioning.

### 2.3 Medial frontal cortex and hippocampal neurons encode Pavlovian unsigned error signals

At the population level, unsigned prediction errors (uPEs, correlating with the state prediction error regressor generated by the model-based learning agent) were decodable at the time of outcome in both dACC and preSMA (both *p* < 0.01, permutation test, Fig. 3A). Additionally, in the encoding analysis of individual neurons, a significant number of neurons coding for uPEs at outcome were present in dACC (12.5%, *p* < 0.01, binomial test), preSMA (12.3%, *p* < 0.001, binomial test), but also hippocampus (10.2%, *p* < 0.05, binomial test, Fig. 3B). In order to determine whether neurons responding to these uPE signals might be better explained by a signed PE signal, we also included a regressor corresponding to the value of the outcome minus the expected value at outcome time, corresponding directly to a signed PE signal. When including a regressor carrying this signal in the same neural analysis from which the uPEs were found, we found no significant evidence for neural encoding of signed PEs in either ACC or preSMA (or elsewhere) (Fig. 3C). Qualitatively similar results were obtained even if generating the signed PEs via a standard temporal difference model (not shown). We summarized the normalized firing rates for uPE coding neurons in dACC, preSMA and hippocampus, (Fig. 3E) and determined that the first times in which preferred vs. non-preferred uPE activity differed in each region were 0.71s, and 0.43s and 0.45s, respectively (*p* < 0.05, t-test). Additionally, at the time of the outcome, we also found a significant proportion of neurons correlated with the outcome itself (reward vs. neutral) in dACC, even when correcting for prediction errors (10.7%, *p* < 0.05, binomial test, Supplementary Figure S2).

**Figure 3:**
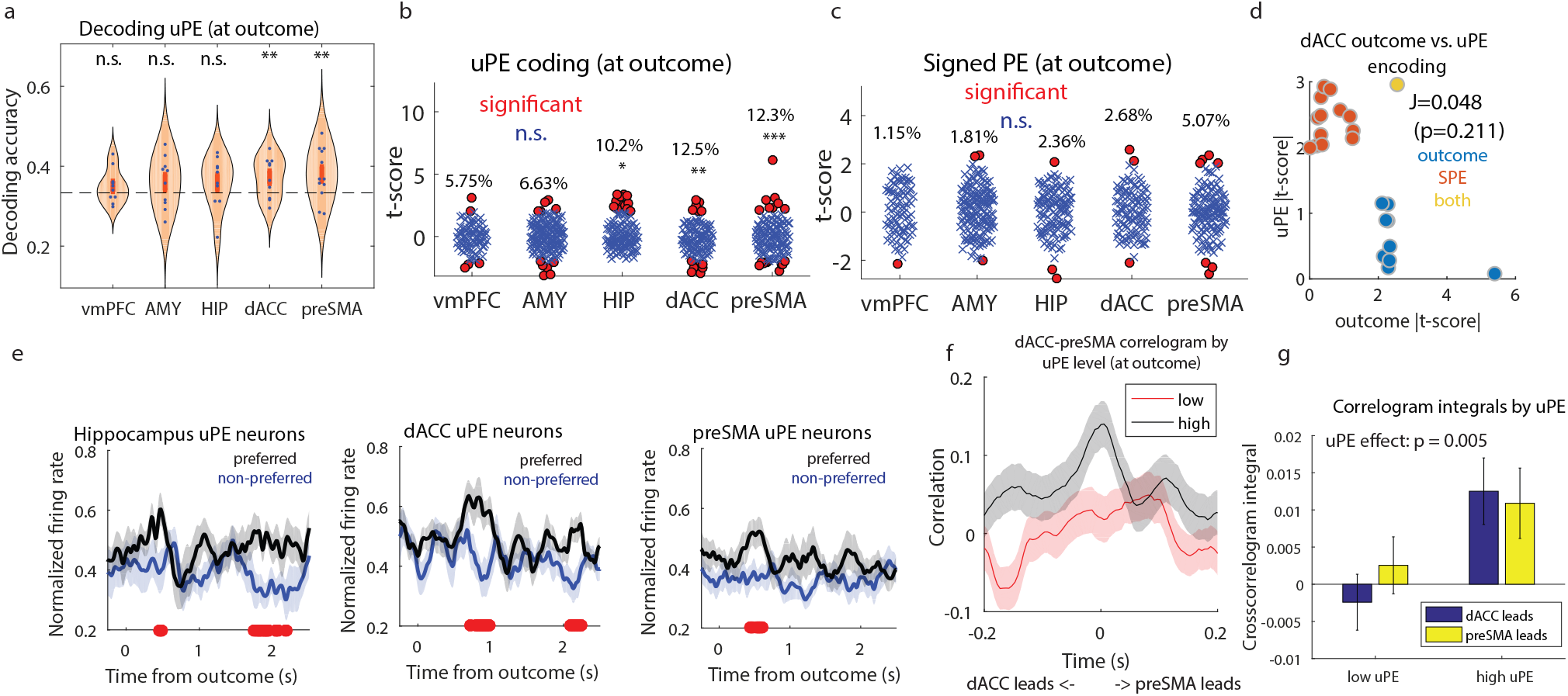
Unsigned prediction error encoding. (a) Decoding accuracy for uPE during outcome presentation. Each dot indicates accuracy in one session, stars indicate significance across sessions with a bootstrapped null distribution, corrected across areas. Bars and dashed lines indicate standard error and chance level, respectively. (b) T-scores for every neuron in each brain area, for a GLM predicting spike counts during outcome presentation with uPE as a regressor. Red dots indicate significant neurons and stars indicate significance across the entire region, corrected across areas. (c) Same, for signed PE. (d) Absolute t-scores for all dACC neurons, comparing neurons coding for outcomes (blue), uPE (orange), or both (yellow). (e) Time course of normalized firing rates in dACC neurons which coded outcomes. Trials are separated between preferred (black) versus non-preferred (blue) outcomes, and red dots indicate time points with a significant difference between trajectories. (f) Spike-spike cross-correlation between dACC and preSMA neurons recorded in the same sessions. Correlograms were computed separately by level of uPE (red: low; black: high). (g) Correlogram integrals by level of uPE (low or high). Integrals were computed as a summary metric, separately in the positive (preSMA leads, yellow) and negative (dACC leads, blue) time lag regions.

### 2.4 Unsigned error signals do not correspond to general state-prediction errors

SPE signals are hypothesized to play a key role in learning of stimulus-stimulus associations (so as to facilitate learning a state-space model necessary for model-based inference). We therefore next set out to test whether the uPE signals described above correspond to state-prediction errors as defined by the model-based agent. A SPE signal should not only signal unexpected states occurring at the time of outcome receipt (where encountering an unexpected outcome can be considered to be an unexpected state), but also during presentations of unexpected proximal stimuli. Because associations between distal and proximal stimuli were varied across blocks, our experimental design elicited variance in state prediction error signals at the time of proximal stimulus delivery. We could therefore test whether the same neurons responding to the uPE signal at the time of outcome were also responding to unexpected stimuli at the time of the proximal stimulus. We tested for neurons correlating with unsigned error signals generated by the model at the time of proximal stimulus delivery. We found no significant neuron count in any brain area (*p* > 0.05 for all brain areas, binomial test), and we only found decoding evidence for a population of neurons coding for SPE-like signals at the time of proximal stimulus in the hippocampus (*p* < 0.05 for all brain areas, permutation test, Supplementary Fig. S1).

Thus, the unsigned prediction error signals we observed in the dACC and preSMA appear to be unique to the time of outcome delivery, suggesting that their response profiles are more specific than would be expected for a general state-prediction error signal. Instead, these signals appear to be specific to salient biologically relevant outcomes (such as visual cues signaling food rewards or their absence) that are unexpected, and respond little to stimuli that have no established value such as the fractal stimuli presented at the proximal stimulus time. In the hippocampus, results were less conclusive, given the positive results in uPE encoding at outcome and uPE decoding at the time of proximal stimulus presentation, followed by null results in uPE decoding at outcome and uPE encoding at proximal time. Such a discrepancy between single neuron encoding results and population decoding results can be the result of a distributed code or the presence of several weakly coding neurons.^55, 56^

### 2.5 Cross-correlations in medial frontal cortex neuronal pairs

Since we found that dACC neurons code for both outcomes and uPEs, we next tested whether the same group of neurons encoded these two variables (Fig. 3D). We found that only one dACC neuron significantly coded both variables, which was not more than expected by chance (p=0.21, Jaccard overlap test). Therefore, coding of outcome and uPE in dACC takes place in largely distinct and dissociable dACC neuronal populations in our dataset.

Given the robust encoding of uPEs in dACC and preSMA, in both single neurons and neural populations, we tested the hypothesis that cross-correlations for dACC-preSMA neuron pairs were modulated by uPEs in the outcome time period. For this, we split all trials into 3 uPE tertiles and defined the first and third tertiles as low and high unsigned error trials, respectively, discarding the middle tertile. Then, for the low and high unsigned error trials, we separately computed spike-spike cross-correlations for all simultaneously recorded dACC-preSMA neuron pairs (see Materials and Methods for details, Fig. 3F). Finally, to summarize our results, we computed cross-correlogram integrals, separately for the positive and negative time lag periods. A peak in the positive time lag region indicates that preSMA spikes preceded dACC spikes more often, whereas a peak in the negative time lag region indicates that dACC spikes preceded preSMA spikes more often.

We performed an ANOVA on the correlogram integrals, with leading region and uPE level as factors, and obtained a significant effect of uPE level (*p* = 0.005, Fig. 3G), but no significant effect for leading region nor an interaction. Taken together, these results shed light on the relationship between preSMA and dACC in prediction error coding during Pavlovian conditioning. In summary, both regions contained predictive information for uPE, and the degree to which dACC and preSMA spikes were aligned in time correlated with the degree of uPE, with no directional preference between the two regions. One possible interpretation of this result is that uPE signals are computed elsewhere in the brain and influence activity in these two areas within a similar time scale.

### 2.6 Expected value coding in vmPFC and amygdala

Next we performed a population decoding analysis for expected values and found significant decoding of proximal EVs in vmPFC and amygdala neural populations (vmPFC: *p* < 0.05; amygdala: *p* < 0.01, permutation test, Fig. 4A). Unlike in population decoding, we did not find a significant neural count for EV coding in any of the brain areas. Nevertheless, given that EV information was present in a distributed code through both neural populations, we tested whether correlations across vmPFC-amygdala neuron pairs were modulated by the level of EV in each trial. Excluding the middle tertile of EV trials, we computed shuffle-corrected correlograms for low and high EV trials (Fig. 4B) and obtained summary correlogram statistics by obtaining their integrals (Fig. 4C).

**Figure 4:**
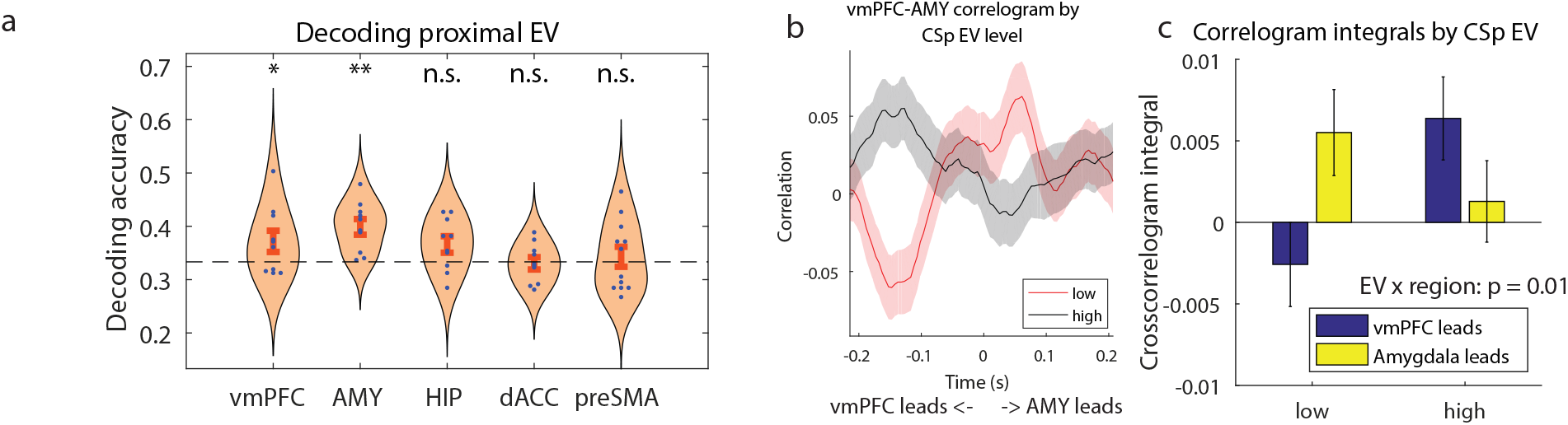
Expected value coding in vmPFC and amygdala. (a) Decoding accuracy for EV proximal stimulus. Each dot indicates accuracy in one session, stars indicate significance across sessions with a bootstrapped null distribution, corrected across areas. Bars and dashed lines indicate standard error and chance level, respectively. (b) Spike-spike cross-correlation between vmPFC and amygdala neurons recorded in the same sessions. Correlograms were computed separately by level of EV (red: low; black: high). (c) Correlogram integrals by level of EV (low or high). Integrals were computed as a summary metric, separately in the positive (amygdala leads, yellow) and negative (vmPFC leads, blue) time lag regions.

Given that vmPFC neurons predictively encoded both stimulus identities and expected values, we next tested whether there was a significant overlap between these two vmPFC neuron subgroups. We found no significant overlap between expected value coding neurons and presumed identity coding neurons in vmPFC (p=0.171, Jaccard overlap test), indicating that predictive identity decoding and outcome decoding in vmPFC are independent subpopulations in this dataset.

We found that the average cross-correlogram peaked at +60ms (amygdala leading) for low EV trials and peaked at -0.127ms (vmPFC leading) for high EV trials, indicating an inversion in the most likely leading region in spike correlations as a function of EV level (Fig. 4B). Additionally, an ANOVA on correlogram integrals, with leading region and EV level as factors revealed a significant interaction between these two factors (*p* = 0.01, Fig. 4C). Taken together, these results shed light on the relationship between amygdala and vmPFC in predictive value coding during Pavlovian conditioning. While both regions contained predictive information for value, which region’s spikes tended to precede the other’s was determined by the expected valence of the outcome. This finding suggests the possibility, in principle, that values learned through Pavlovian conditioning have the ability to dynamically alter connectivity patterns between amygdala and vmPFC, regions which have been shown to selectively code value-based cues.^37, 42^

## 3 Discussion

In this study, we probed for neural representations of associations and learning signals in the human brain during Pavlovian conditioning at the level of single neurons. We were especially focused on determining whether we could find evidence for prospective coding of information about the identity of upcoming events during Pavlovian conditioning. We hypothesized that neurons in the ventromedial prefrontal cortex would be involved in such prospective identity coding, on account of prior evidence about a role for rodent orbitofrontal cortex neurons in identity coding, as well as fMRI results from human orbitofrontal cortex.^25, 37, 38, 41, 42^ We found that neurons in the human vmPFC did indeed encode information about subsequent stimuli that are expected to occur, suggesting that neurons in this area are involved in encoding stimulus-stimulus relationships. However, we did not find evidence for a comparable encoding of predictive stimulus information in the amygdala, another structure previously implicated in model-based Pavlovian conditioning in humans.^57^ The finding that vmPFC neurons (including a subset of neurons in the medial orbitofrontal cortex) are involved in encoding stimulus-stimulus associations is compatible with a theoretical proposal implicating the orbitofrontal cortex in facilitating a cognitive map of task contingencies.^58^ This theoretical account especially implicates the orbitofrontal cortex in making inferences about hidden states. In the present study we tested for neurons encoding representations of the most likely stimulus to be subsequently presented given the current associative structure, which could be considered a relatively simple form of inference over states. This type of stimulus-stimulus identity coding during Pavlovian conditioning could form part of the neural substrates for model-based Pavlovian conditioning.^4, 25, 57^

We also aimed to test for neurons coding for state prediction errors, a signal that could theoretically be deployed by a model-based agent to underpin learning of stimulus-stimulus associations so as to build a cognitive model of the associative structure needed for model-based Pavlovian inference. We found individual neurons and population level encoding of an unsigned error signal in the hippocampus, preSMA and dACC at the time of outcome receipt. However, a state prediction error signal should also respond to unexpected stimuli even if they do not have strong affective significance, such as the fractal stimuli presented in our sequential Pavlovian paradigm. An unexpected presentation or omission of a particular stimulus in the sequential chain should elicit a state prediction error response. While we found robust state-prediction error responses at the time of outcome delivery, these same neurons did not respond to unexpected stimulus presentations or omissions, nor did we find evidence for a separate population of neurons responding to unexpected non-outcome stimuli. Thus, the unsigned error signal we found appears not to meet the criteria necessary to be deemed a state prediction error as defined by the theoretical model. We also ruled out other explanations for these unsigned prediction error signals at the time of outcome such as that they are signed reward prediction errors or merely signaling the valence of outcome delivery. We found little evidence for signed reward prediction errors in these regions, and neurons responding to the unsigned error signals could be clearly dissociated from a separate population of neurons in dACC responding instead to outcome valence. These outcome-specific unsigned prediction neurons appear to exhibit similar properties to a subset of dopaminergic neurons found to respond to unexpected outcomes irrespective of their valence.^59^ One possible explanation for these error signals is that they play a specific role in mediating learning of stimulus-outcome associations where outcome properties are features of outcomes other than valence such as outcome identity, but they are not involved in more general processes pertaining to learning of a cognitive map.

Intriguingly, we found these unsigned outcome-specific error signals in the hippocampus as well as in dorsomedial prefrontal cortex. The hippocampal error signals we observed appeared to be qualitatively different to those in dorsomedial prefrontal cortex. Whereas in the preSMA and dACC some neurons increased their firing rate in response to an increase in unsigned error while others decreased their firing rate in response to an increase in unsigned error, the hippocampal neurons were found to be exclusively positively correlated with unsigned error (Fig. 3B, all t-scores over 0), such that activity in these neurons only increased as a function of unsigned error. The functional significance of this finding is unclear, but it merits attention in future studies so as to determine its robustness and significance. It is also notable that we could also decode error signals at the time of proximal stimulus presentation relating to how unexpected that particular stimulus was, albeit this result should be viewed as tentative given it was only weakly significant. The finding of a role for hippocampal neurons in encoding errors associated with learning of stimulus-outcome relationships and potentially even in stimulus-stimulus associations, adds further insight into the putative role of the hippocampus in Pavlovian conditioning, as well as in model-based inference more generally.^28, 47, 48^ One hypothesis is that these hippocampal error neurons play a general role in facilitating predictive learning about stimulus-stimulus associations as part of a cognitive map, even as they are encountered passively through Pavlovian learning.

Our findings also shed unique light on the functions of dorsomedial prefrontal cortex in error monitoring. There is considerable evidence to suggest that neurons in the dorsomedial prefrontal cortex are involved in error signaling during or shortly after the active performance of motor responses, involved in particular in signaling when an incorrect response has occurred.^49–51, 53^ Crucially, the present study reveals that neurons in dmPFC also track error signals even when no motor responses are being actively performed by the participant to obtain the outcome, because in Pavlovian conditioning, unlike instrumental conditioning, the organism passively observes the relationship between stimuli and outcomes. The unsigned error signals we observed here are thus not in any way related to motor or response errors. A number of theories of dmPFC function in error monitoring have instead emphasized a role for the reward prediction error signal carried by the dopaminergic system in driving error-related activity changes in these areas, especially the error-related negativity detected in scalp EEG.^60–63^ Such a reward error signal would signal a deviation in expected reward obtained, as opposed to depicting errors in responses performed per se. Interestingly, in the present study, the unsigned prediction error we observed in dACC and preSMA neurons was clearly distinct from signed PEs of the sort associated with phasic dopaminergic activity. Unlike the classic signed reward prediction error signal, these neurons increased in activity at the time of outcome for any outcome that was unexpected irrespective of whether it was an unexpected reward outcome or an unexpected omission of reward. Moreover, these unsigned error signals were detected even after adjusting for the effects of signed reward prediction errors in the same statistical model. These findings suggest broader functions of dmPFC neurons in error signaling beyond either response-related error monitoring or signed reward prediction error signaling.

In addition to finding neuronal signals related to identity coding and unsigned error coding that are orthogonal to value, we also found evidence of a role for neurons in the vmPFC and amygdala in encoding stimulus-value at the population level. These brain regions are deeply interconnected functionally and anatomically.^64^ For instance, primate OFC displays strong anatomical projections to amygdala,^65^ and vice-versa,^66^ and the same is true for vmPFC proper.^67^ Importantly, neurons in OFC and amygdala have been shown to respond selectively in anticipation of rewarding or aversive outcomes in rodents and monkeys,^37, 42^ while BOLD responses in humans have been found to correlate with expected value across an array of paradigms including Pavlovian appetitive conditioning.^6, 13, 25, 40, 68, 69^ Above and beyond its aforementioned role in predictive value coding in tandem with OFC, the amygdala has been shown to track expected values during decision making,^38–40^, ^43–45, 55^ while lesion studies have suggested a causal role for this brain area in utilizing learned expected values for guiding behavior.^70–72^ Additionally, targeted amygdala lesions in rodents have been shown to abolish motivational or identity-specific effects of Pavlovian cues on operant behavior.^73^

Importantly, recent opto- and chemogenetic circuit dissection studies in rodents have demonstrated that bidirectional amygdala-OFC interactions are critical for both encoding and retrieval of reward value, and for the construction of cue-triggered reward expectations.^74, 75^ Our findings provide preliminary evidence in support of a similar network-level substrate in humans. We investigated whether spiking activity in simultaneously recorded vmPFC and amygdala neuron pairs correlated in time, and found that cross-correlation patterns were modulated by the expected value of the conditioned stimulus. Specifically, amygdala neuron spikes tended to precede vmPFC spikes when patients were exposed to a cue which predicted a low value outcome, while the opposite occurred for high value cues. These findings provide further insight into the nature of the interactions between these regions, and are congruent with prior literature suggesting a critical role for interactions between the amygdala and OFC in value computations. For instance, a lack of healthy amygdala input induces significant changes to vmPFC activity during reward learning and decision making^76^ and impairs the formation of neural ensembles in OFC to represent new contingencies during reversal learning.^77^ When taken together with this prior literature, our results are compatible with converging evidence positioning amygdala as a center for predictive value coding acting in consonance with OFC, which creates associations between stimulus identities and outcomes.

Broadly, our findings provide a glimpse into how Pavlovian conditioning takes place in human prefrontal cortex neurons as well as in the amygdala and hippocampus, adding to an electrophysiology literature in other species as well as human fMRI studies. We provide unique evidence that human vmPFC neurons encode representations of predictive identity, consistent with previous findings from BOLD fMRI and electrophysiological results in rodents. Moreover, we found evidence that neurons in dorsomedial prefrontal cortex such as the preSMA and dACC and the hippocampus encode error signals related to unexpected outcomes that could be used to facilitate learning of stimulus-outcome associations involving features other than value.

## Supplementary Material for

## 1 Materials and Methods

### 1.1 Electrophysiology and recording

We used Behnke-Fried hybrid depth electrodes (AdTech Medical), positioned exclusively according to clinical criteria. Broadband extracellular recordings were performed with a sampling rate of 32 kHz and a bandpass of 0.1-9000Hz (ATLAS system, Neuralynx Inc.). The data set reported here was obtained bilaterally from hippocampus (HIP), amygdala (AMY), ventromedial prefrontal cortex (vmPFC), dorsal anterior cingulate cortex (dACC), and pre-supplementary motor area (preSMA) with one macroelectrode on each side. Each macroelectrode contained eight 40 *µ*m diameter microwires. Recordings were locally referenced, with one of the eight microwires serving as a reference in every bundle.

### 1.2 Patients

Twelve patients (eight females) were implanted with depth electrodes for seizure monitoring prior to potential surgery for treatment of drug resistant epilepsy. One of the patients performed the task twice, totalling 13 recorded sessions. Human research experimental protocols were approved by the Institutional Review Boards of the California Institute of Technology and the Cedars-Sinai Medical Center. Electrode location was determined based on preoperative structural MRI and postoperative MRI and/or CT scans obtained for each patient as described previously Minxha et al. (2020). Patient information (sex and age) is displayed in Supplementary Table 1.

### 1.3 Pavlovian conditioning task

Patients performed a sequential Pavlovian conditioning task Pauli et al. (2015) in which two fractals were presented in sequence, acting as distal and proximal conditioned stimuli (CSd, CSp, respectively). Following the CS pair, a video outcome was presented, either in the form of a hand depositing the patient’s candy of choice into a paper bag, for a positive outcome, or in the form of an empty hand approaching a paper bag, for a neutral outcome. Patients were informed that each time they received a positive outcome contributed to a grand total of actual candy pieces they would receive at the end of the experiment. Every patient experienced exactly 48 win trials and 48 no-win trials. Prior to the task, patients visually inspected and chose one among five possible candy options (Reese’s Pieces, Hershey’s Kisses, York Peppermint, Werther’s Caramel, or Hi-Chew) to be delivered, according to their preference. After the end of the session, they received the closest possible equivalent to 200 calories in their candy of choice, to reward them for the 48 win trials they experienced.

Trial structure is detailed in Fig. 1A. After a 1s fixation period, a distal CS was presented for 3s, followed by a 1s fixation period. Then, a proximal CS was displayed for 3s, followed by an outcome presentation video edited to a length of 3s. Intertrial intervals were jittered between 0.5s-1s. The distal CS was always presented along the vertical axis, either above or below the center of the screen, at a random position. Similarly, the proximal CS was always presented in a random position along the horizontal axis, inside one of two possible squares positioned to the left or the right of the screen. As an attention check, patients were asked to perform a button press whenever they saw a CSp inside one of the squares, reporting which of the two squares it was. To ensure this remained a purely Pavlovian paradigm, patients were instructed that these button presses did not affect the outcome of the trial in any way.

Each session contained a total of 96 trials, split into 4 blocks of 24 trials, with a two minute break between blocks. Within each block, there were 4 possible CS, 2 distal (A,B) and 2 proximal (X,Y). The identity of the distal stimuli was re-selected in every block, but the two proximal stimuli were kept the same throughout the entire session, though their valences were reversed between blocks. There were four trial types in total (Fig. 1C): two common transitions (A-X-Reward and B-Y-No reward) and two rare transitions (A-Y-No reward and B-X-Reward), meaning that the CSp was always fully predictive of the outcome, while the CSd was only partially predictive. Trial frequencies were 37.5% for each of the common types and 12.5% for each of the rare types. Every block always had exactly 9 trials of each common type and 3 trials of each rare type. Trial type order was selected randomly, except for the first 4 trials of each block, which we enforced to be of the common type, selected randomly.

The 10 fractals used in each session (2 CSd fractals for each block and 2 CSp fractals for the whole session) were chosen specifically for each patient out of a set of 24 possible fractals using the following procedure: before the beginning of the task, patients rated all 24 fractals for their subjective preference using a sliding bar from extremely unpleasant to extremely pleasant. The 10 stimuli which elicited the most neutral responses were selected for the experiment and assigned randomly to their CSd or CSp roles. All the 24 fractals had the same mean luminance. We enforced this by enforcing equal average RGB values across pixels for all fractals, along the (0,0,0) to (255,255,255) spectrum. At the end of every block, patients rated the 4 fractals they experienced for how pleasant they were. We z-scored ratings for each patient and used aggregate stimulus rating changes across patients as a metric of Pavlovian conditioning.

### 1.4 Eye tracking

We tracked patients’ pupil diameter during the task as a candidate conditioning metric Pauli et al. (2015, 2019), Pool et al. (2019). We used a EyeLink 1000 system (SR Research Ltd.) attached to the bottom of the task screen as described in Minxha et al. (2017), Wang et al. (2019). We calibrated the system using EyeLink’s five point calibration before the session and between blocks (average calibration error = 0.40° *±* 0.17). Patients were instructed to fixate during fixation periods but were free to gaze anywhere in the screen otherwise. Pupil data was preprocessed to remove blinks and outlier points further than 5 s.d. from the mean diameter. We interpolated missing values removed in this way with the closest previous value and then filtered data with a 50ms moving average window. Pupil diameters were normalized in every trial relative to the initial 1s fixation period, using the average diameter in that period as a baseline. Statistical analyses were then performed using average diameter changes in the 0.5s-3s time windows after CSd and CSp presentations.

### 1.5 Computational model of learning

We used a normative model-based model to obtain estimates of how patients encoded transition probabilities between task states, stimulus expected values, and state prediction errors. We adapted a model used for a sequential instrumental task Gläscher et al. (2010), to estimate a matrix *T* (*s, s*′) for the transition probabilities from start states *s* to end states *s*′. Given distal states (A,B), proximal states (X,Y), and outcome states (reward, R; no reward, N), we defined the transition matrix T with transition probabilities *t*_*ss*_′ as:

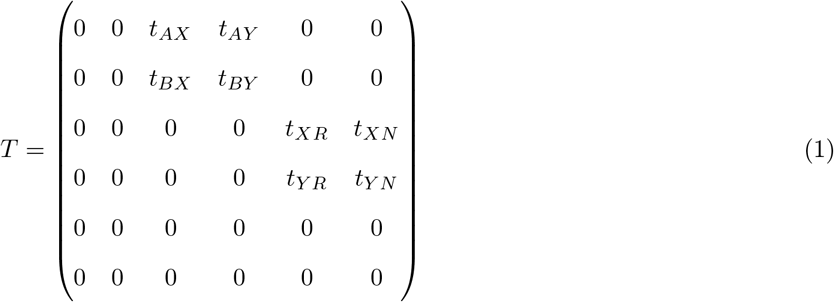

The *t* values were initialized to 0.5 and the remaining values in the matrix were chosen to be 0, reflecting the constraints of the task’s transition structure. At each step, after transitioning from some state *s* to an end state *s*′, a state prediction error is computed according to the following equation:

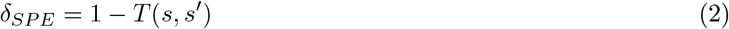

The transition matrix *T* is then updated using a learning rate *η*, chosen normatively to be 0.22, adopting a value from a previously validated model-free model Pauli et al. (2015).

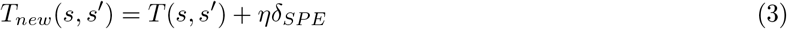

All the other values in T for states not transitioned into were also updated with *T*_*new*_(*s, s*″) = *T* (*s, s*″)(1 *− η*) to ensure each the sum of each row of T stays equal to 1, as it should reflect total probability.

Finally, the value of each state could be computed assuming the values of arriving into the reward and no reward states were 1 and 0, respectively:

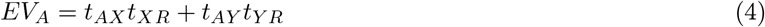

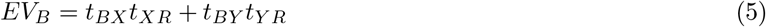

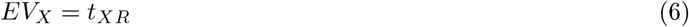

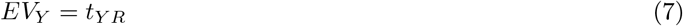

Since we were interested in tracking neural responses to stimulus identity associations, we used the inferred transition matrix T to determine the identity of the most likely proximal stimulus to follow distal stimulus presentation. We refer to this identity as *CSp presumed identity*.

We chose a normative model-based learning mechanism for analyzing the data because this model allowed us to estimate the presumed identity of subsequent stimuli in the associative chain, thereby allowing us to probe stimulus-stimulus mappings in the task. It is important to note that a model-free temporal difference (TD) model (cite Sutton) could be used instead to model this data, and that such a model generates highly similar time-series for value signals as we used in the behavioral and neural analyses (though it would not allow us to make predictions about stimulus identities). Using such a TD-like model, a signed reward prediction error can be estimated, and the absolute value (unsigned) of this error signal also strongly correlates with SPE signals used in the analysis. While we implemented such a TD model alongside the MB model and used it to analyze the data, we do not report on the results of this analysisbecause it produces highly similar results about value encoding as reported using thr existing MB model, and because this TD model does not allow us to test for stimulus identity representations.

### 1.6 Neural data pre-processing

We performed spike detection and sorting with the semiautomatic template-matching algorithm OSort Rutishauser et al. (2006). Across all 13 sessions, we obtained 86 vmPFC, 165 amygdala, 119 hippocampus, 137 preSMA and 103 dACC putative single units (610 total). We refer to these isolated putative single units as “neuron” and “cell” interchangeably.

### 1.7 Single neuron encoding analysis

To quantify how single neuron activity correlated with several variables of interest, we performed a Poisson generalized linear model (GLM) analysis. As a dependent variable, we measured spike counts in three time windows: CSd window, from 0.25s to 3s after CSd presentation; CSp window, from 0.25s to 3s after CSp presentation; outcome window, from 0.25s to 2.25s after outcome presentation. Table 1 includes all dependent variables and the time windows in which they were tested. We defined RPEs as the difference between proximal EV at the time of outcome and the outcome valence itself.

**Table 1:**
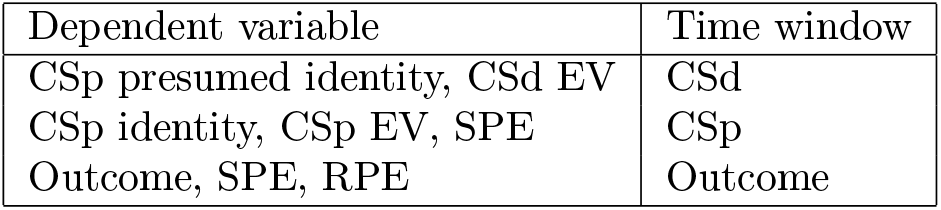
Dependent variables and respective time windows for Poisson GLM and population decoding analyses.

### 1.8 Jaccard index test

After performing Poisson GLM encoding analyses, we tested whether the sub-populations of neurons which were sensitive to two variables of interest had significant overlap. For this, we computed the Jaccard index Jaccard (1912) of overlap between neurons sensitive to each of the variables *X* and *Y*, where *N*_*X*_ and *N*_*Y*_ indicate the number of neurons sensitive to the variables X and Y, respectively, and *N*_*X,Y*_ indicates the number of neurons concurrently sensitive to both variables:

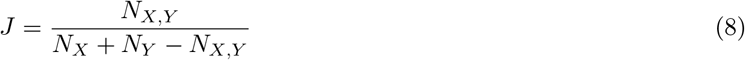

To compute p-values for each comparison between two variables, we bootstrapped a null distribution of Jaccard indexes using 1000 reshuffles, considering X and Y are independent variables with a false positive rate of *p* = 0.05.

### 1.9 Population decoding analysis

We performed population decoding analyses by training a linear support vector machine (SVM) with MATLAB’s function *fitcsvm*. In each session, we defined a population activity matrix X of dimensions (*nTrials, nNeurons*) by counting spikes in each trial within the same time windows from the encoding analysis. We then defined the decoded variable *y* as the same dependent variables from Table 1. To reduce the decoding problem to a classification task, we binned the continuous regressors (EV, SPE) into 3 tertiles before training the SVM with MATLAB’s multiclass function *fitcecoc*. Cross-validation was performed by training on 2 trial blocks and testing on the 2 remaining trial blocks of each session. Since there were 4 blocks in total, we repeated the procedure for every possible combination of train/test blocks, resulting in 6 cross-validation folds. Test accuracies were averaged across folds and reported separately for each session. To obtain test accuracy significance levels, we repeated this entire procedure 500 times while shuffling the decoded variable *y* and compare the null mean test accuracy obtained in this manner with the true test accuracy, for each brain region. Finally, we corrected significance thresholds for the number of tested brain areas. For this analysis, we excluded one session, which contained only 1 block of trials.

### 10.10 Cross-correlation analysis

To measure how neural activity across brain areas of interest was correlated, we performed a spike-spike crosscorrelation analysis, with shuffle correction Brody (1999). First, we divided all trials in EV tertiles, excluding the middle tertile to obtain low EV and high EV trial groups. Then, for each trial group, we computed cross-correlations for each neuron pair containing neurons A and B recorded from the same session, but different brain areas (e.g. each neuron pair contained one vmPFC neuron vs. one amygdala neuron).

For two spike trains 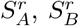 (binned at 50ms with 5ms steps, constrained to CSp presentation time window), recorded from neurons A and B on trial *r*, we define the cross-correlogram of each trial as:

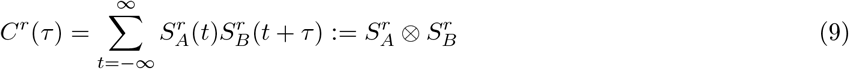

Defining the <> operator as averaging across trials, we defined the shuffle-corrected cross-correlogram as:

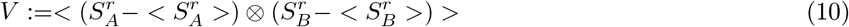

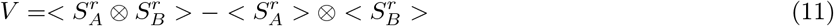

The shuffle-correction procedure corrects for time-locked co-variation that might be caused to both neurons concurrently by stimulus presentation. Additionally, it ensures that the expected value of *V* is 0 if *S*_*A*_ and *S*_*B*_ are independent.

Finally, the reported shuffle-corrected correlation results were obtained by averaging the *V* vectors across all neuron pairs. To summarize the correlation results, we obtained the integrals from the positive and negative time lag regions and performed an ANOVA with EV level (low vs. high) and time lag sign (which region leads) as factors.

## 2 Pavlovian outcome coding

At the time of the outcome, we found a significant proportion of neurons correlated with outcome (reward vs. neutral) in dACC (10.7%, *p* < 0.05, binomial test, Supplementary Fig. 2A). An example dACC neuron which coded outcomes is shown in Supplementary Fig. 2B. Additionally, for outcome encoding dACC neurons, we tested when normalized firing rates differed for preferred versus non-preferred outcomes, and found that they first differed 1.08s after outcome onset (*p* < 0.05, t-test, uncorrected). Out of all significant outcome dACC neurons, 10*/*12 had negative t-scores, indicating they fired less in trials with positive outcomes. This is more than expected by chance assuming an equal probability of positive and negative t-scores (*p* = 0.003, binomial test). At the population level, we found significant decoding of outcomes in dACC (*p* < 0.001, permutation test,Supplementary Fig. 2C). However, above and beyond the single neuron encoding results, outcome was also decodable from the hippocampus (*p* < 0.05, permutation test) and preSMA (*p* < 0.05, permutation test), highlighting a widespread population-level representation of outcome valence. One possible interpretation is that weakly coding neurons provided a distributed code for discriminating between positive and neutral outcomes when considered jointly (Stefanini et al. 2020). These results are consistent with previous findings on tracking feedback in medial frontal cortex neurons (Fu et al. 2019, Aquino et al. 2021).

## Supplementary Figures

**Figure 1:**
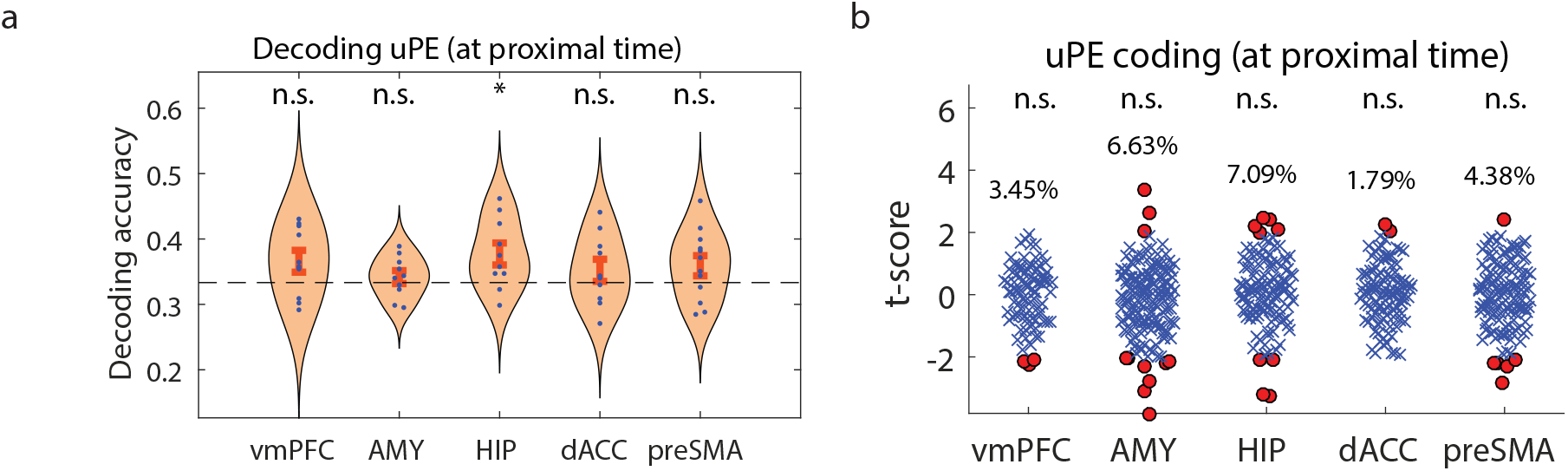
Unsigned prediction error encoding at proximal time. (a) Decoding accuracy for uPE during proximal stimulus presentation. Each dot indicates accuracy in one session, stars indicate significance across sessions with a bootstrapped null distribution, corrected across areas. Bars and dashed lines indicate standard error and chance level, respectively. (b) T-scores for every neuron in each brain area, for a GLM predicting spike counts during proximal stimulus presentation with uPE as a regressor. Red dots indicate significant neurons and stars indicate significance across the entire region, corrected across areas.

**Figure 2:**
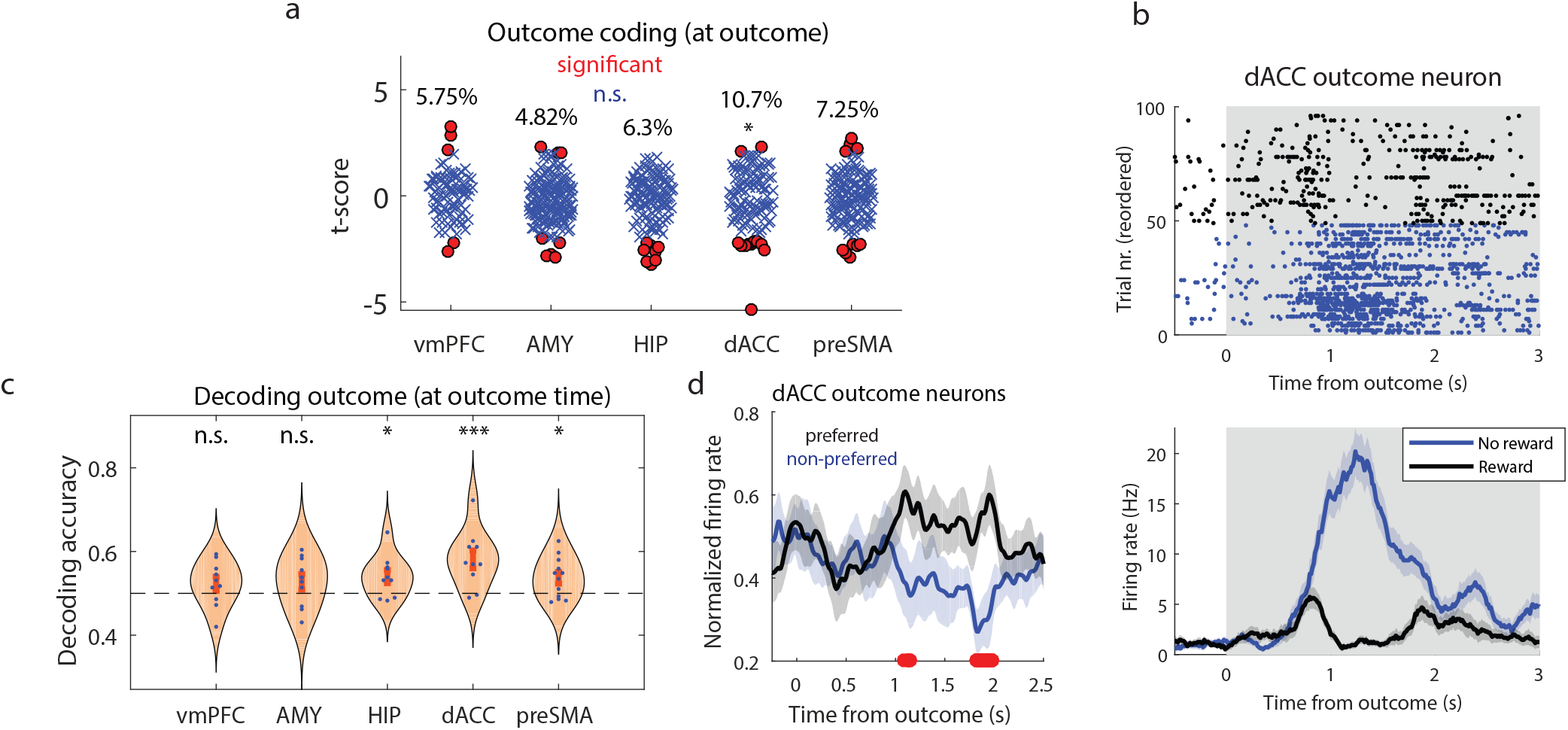
Outcome encoding in dACC. (a) T-scores for every neuron in each brain area, for a GLM predicting spike counts during outcome presentation with outcome as a regressor. Red dots indicate significant neurons and stars indicate significance across the entire region, corrected across areas. (b) dACC neuron whose activity during correlates with outcome (blue: no reward; black: reward). Top: raster plot; Bottom: PSTH. (c) Decoding accuracy for outcome during outcome presentation. Each dot indicates accuracy in one session, stars indicate significance across sessions with a bootstrapped null distribution, corrected across areas. Bars and dashed lines indicate standard error and chance level, respectively. (d) Time course of normalized firing rates in dACC neurons which coded outcomes. Trials are separated between preferred (black) versus non-preferred (blue) outcomes, and red dots indicate time points with a significant difference between trajectories.

## Supplementary Tables

**Table 2:**
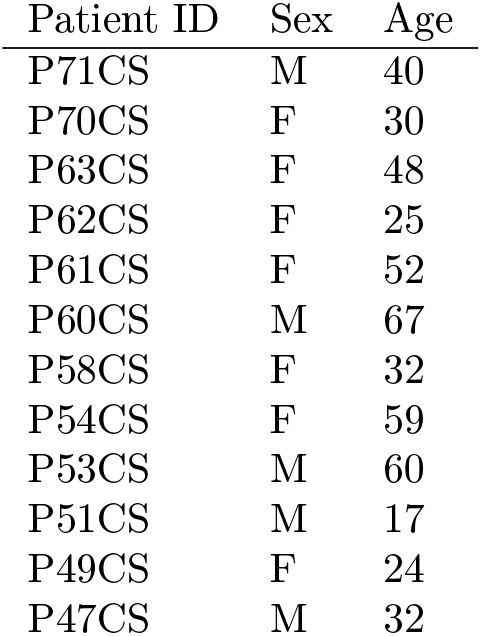
Patient labels, sex, and age for all subjects.

