## Supplementary Information for "Encoding of predictive associations in human prefrontal and medial temporal neurons during Pavlovian conditioning"

$$T = \begin{pmatrix} 0 & 0 & t_{AX} & t_{AY} & 0 & 0 \\ 0 & 0 & t_{BX} & t_{BY} & 0 & 0 \\ 0 & 0 & 0 & 0 & t_{XR} & t_{XN} \\ 0 & 0 & 0 & 0 & t_{YR} & t_{YN} \\ 0 & 0 & 0 & 0 & 0 & 0 \\ 0 & 0 & 0 & 0 & 0 & 0 \end{pmatrix} \quad (1)$$

The $t$ values were initialized to 0.5 and the remaining values in the matrix were chosen to be 0, reflecting the constraints of the task's transition structure. At each step, after transitioning from some state $s$ to an end state $s'$, a state prediction error is computed according to the following equation:

$$\delta_{SPE} = 1 - T(s, s') \quad (2)$$

The transition matrix $T$ is then updated using a learning rate $\eta$, chosen normatively to be 0.22, adopting a value from a previously validated model-free model Pauli et al. (2015).

$$T_{new}(s, s') = T(s, s') + \eta\delta_{SPE} \quad (3)$$

All the other values in T for states not transitioned into were also updated with $T_{new}(s, s'') = T(s, s'')(1 - \eta)$ to ensure each the sum of each row of T stays equal to 1, as it should reflect total probability.

Finally, the value of each state could be computed assuming the values of arriving into the reward and no reward states were 1 and 0, respectively:

$$EV_A = t_{AX}t_{XR} + t_{AY}t_{YR} \quad (4)$$

$$EV_B = t_{BX}t_{XR} + t_{BY}t_{YR} \quad (5)$$

$$EV_X = t_{XR} \quad (6)$$

$$EV_Y = t_{YR} \tag{7}$$

Since we were interested in tracking neural responses to stimulus identity associations, we used the inferred transition matrix T to determine the identity of the most likely proximal stimulus to follow distal stimulus presentation. We refer to this identity as *CSp presumed identity*.

| Dependent variable | Time window |
|---|---|
| CSp presumed identity, CSd EV | CSd |
| CSp identity, CSp EV, SPE | CSp |
| Outcome, SPE, RPE | Outcome |

Table 1: Dependent variables and respective time windows for Poisson GLM and population decoding analyses.

For two spike trains $S_A^r$, $S_B^r$ (binned at 50ms with 5ms steps, constrained to CSp presentation time window), recorded from neurons A and B on trial $r$, we define the cross-correlogram of each trial as:

$$C^r(\tau) = \sum_{t=-\infty}^{\infty} S_A^r(t) S_B^r(t+\tau) := S_A^r \otimes S_B^r \quad (9)$$

Defining the <> operator as averaging across trials, we defined the shuffle-corrected cross-correlogram as:

$$V :=< (S_A^r - < S_A^r >) \otimes (S_B^r - < S_B^r >) > \quad (10)$$

$$V =< S_A^r \otimes S_B^r > - < S_A^r > \otimes < S_B^r > \quad (11)$$

The shuffle-correction procedure corrects for time-locked co-variation that might be caused to both neurons concurrently by stimulus presentation. Additionally, it ensures that the expected value of $V$ is 0 if $S_A$ and $S_B$ are independent.

Finally, the reported shuffle-corrected correlation results were obtained by averaging the $V$ vectors across all neuron pairs. To summarize the correlation results, we obtained the integrals from the positive and negative time lag regions and performed an ANOVA with EV level (low vs. high) and time lag sign (which region leads) as factors.

### 2 Pavlovian outcome coding

At the time of the outcome, we found a significant proportion of neurons correlated with outcome (reward vs. neutral) in dACC (10.7%, $p < 0.05$, binomial test, Supplementary Fig. 2A). An example dACC neuron which coded outcomes is shown in Supplementary Fig. 2B. Additionally, for outcome encoding dACC neurons, we tested when normalized firing rates differed for preferred versus non-preferred outcomes, and found that they first differed 1.08s after outcome onset ($p < 0.05$, t-test, uncorrected). Out of all significant outcome dACC neurons, 10/12 had negative t-scores, indicating they fired less in trials with positive outcomes. This is more than expected by chance assuming an equal probability of positive and negative t-scores ($p = 0.003$, binomial test). At the population level, we found significant decoding of outcomes in dACC ($p < 0.001$, permutation test,Supplementary Fig. 2C). However, above and beyond the single neuron encoding results, outcome was also decodable from the hippocampus ($p < 0.05$, permutation test) and preSMA ($p < 0.05$, permutation test), highlighting a widespread population-level representation of outcome valence. One possible interpretation is that weakly coding neurons provided a distributed code for discriminating between positive and neutral outcomes when considered jointly (Stefanini et al. 2020). These results are consistent with previous findings on tracking feedback in medial frontal cortex neurons (Fu et al. 2019, Aquino et al. 2021).

### Supplementary Figures

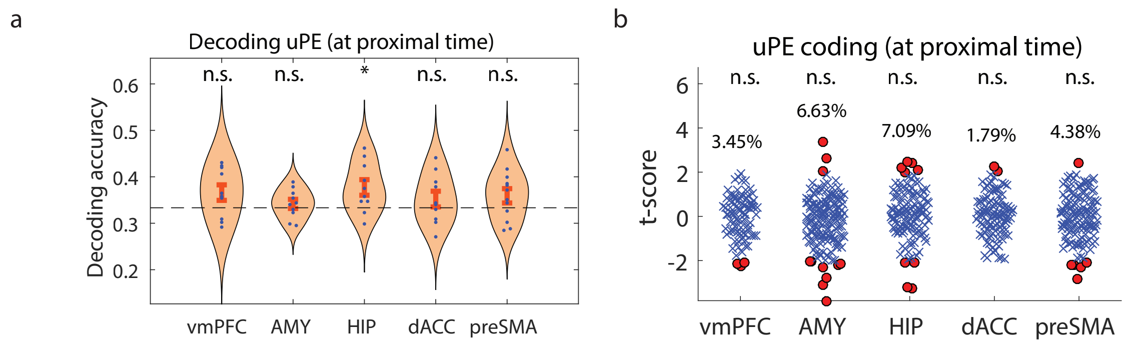


Figure 1: Unsigned prediction error encoding at proximal time. (a) Decoding accuracy for uPE during proximal stimulus presentation. Each dot indicates accuracy in one session, stars indicate significance across sessions with a bootstrapped null distribution, corrected across areas. Bars and dashed lines indicate standard error and chance level, respectively. (b) T-scores for every neuron in each brain area, for a GLM predicting spike counts during proximal stimulus presentation with uPE as a regressor. Red dots indicate significant neurons and stars indicate significance across the entire region, corrected across areas.

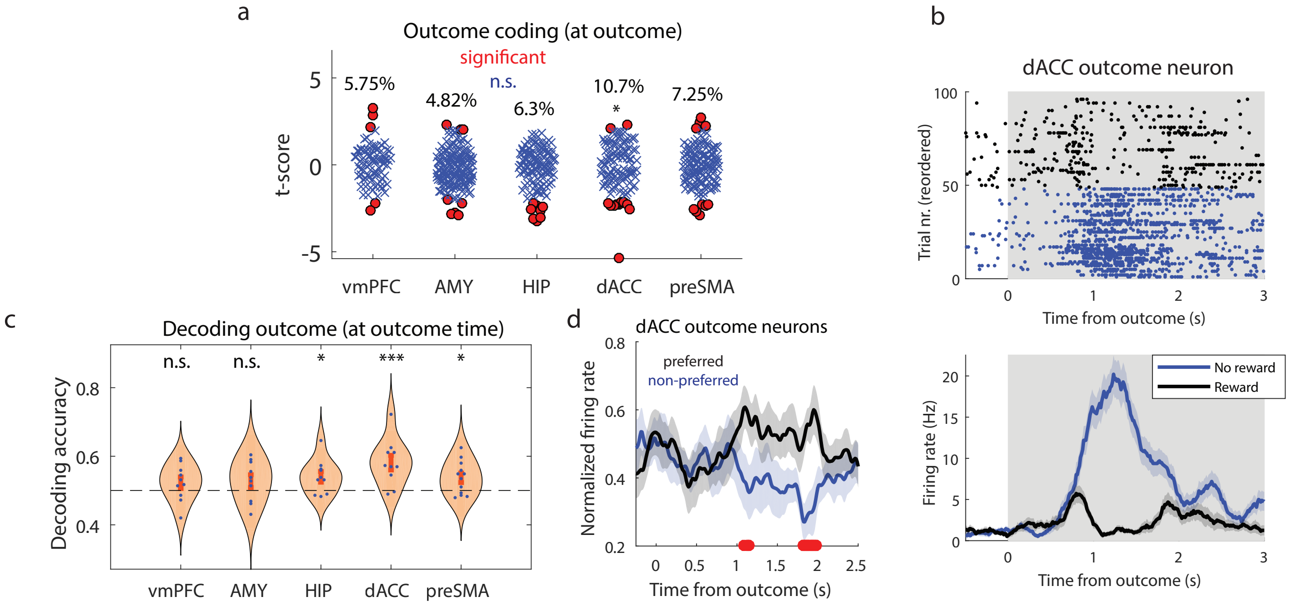


Figure 2: Outcome encoding in dACC. (a) T-scores for every neuron in each brain area, for a GLM predicting spike counts during outcome presentation with outcome as a regressor. Red dots indicate significant neurons and stars indicate significance across the entire region, corrected across areas. (b) dACC neuron whose activity during correlates with outcome (blue: no reward; black: reward). Top: raster plot; Bottom: PSTH. (c) Decoding accuracy for outcome during outcome presentation. Each dot indicates accuracy in one session, stars indicate significance across sessions with a bootstrapped null distribution, corrected across areas. Bars and dashed lines indicate standard error and chance level, respectively. (d) Time course of normalized firing rates in dACC neurons which coded outcomes. Trials are separated between preferred (black) versus non-preferred (blue) outcomes, and red dots indicate time points with a significant difference between trajectories.

### Supplementary Tables

| Patient ID | Sex | Age |
|---|---|---|
| P71CS | M | 40 |
| P70CS | F | 30 |
| P63CS | F | 48 |
| P62CS | F | 25 |
| P61CS | F | 52 |
| P60CS | M | 67 |
| P58CS | F | 32 |
| P54CS | F | 59 |
| P53CS | M | 60 |
| P51CS | M | 17 |
| P49CS | F | 24 |
| P47CS | M | 32 |

Table 2: Patient labels, sex, and age for all subjects.
